# Ca^2+^-activation kinetics modulate successive puff/spark amplitude, duration and inter-event-interval correlations in a Langevin model of stochastic Ca^2+^ release

**DOI:** 10.1101/014431

**Authors:** Xiao Wang, Yan Hao, Seth H. Weinberg, Gregory D. Smith

## Abstract

Through theoretical analysis of the statistics of stochastic calcium (Ca^2+^) release (i.e., the amplitude, duration and inter-event interval of simulated Ca^2+^ puffs and sparks), we show that a Langevin description of the collective gating of Ca^2+^ channels may be a good approximation to the corresponding Markov chain model when the number of Ca^2+^ channels per Ca^2+^ release unit (CaRU) is in the physiological range. The Langevin description of stochastic Ca^2+^ release facilitates our investigation of correlations between successive puff/spark amplitudes, durations and inter-spark intervals, and how such puff/spark statistics depend on the number of channels per release site and the kinetics of Ca^2+^-mediated inactivation of open channels. When Ca^2+^ inactivation/de-inactivation rates are intermediate—i.e., the termination of Ca^2+^ puff/sparks is caused by the recruitment of inactivated channels—the correlation between successive puff/spark amplitudes is negative, while the correlations between puff/spark amplitudes and the duration of the preceding or subsequent inter-spark interval are positive. These correlations are significantly reduced when inactivation/deinactivation rates are extreme (slow or fast) and puff/sparks terminate via stochastic attrition.

## 1 Introduction

Intracellular Ca^2+^ elevations known as Ca^2+^ puffs and sparks (Cheng et al., 1993; Yao et al., 1995) arise from the cooperative activity of inositol 1,4,5-trisphosphate receptors (IP_3_Rs) and ryanodine receptors (RyRs) that are clustered in Ca^2+^ release units (CaRUs) on the endoplasmic reticulum/sarcoplasmic reticulum (ER/SR) membrane (see Berridge (1993); Bers (2002) for review). Single-channel Ca^2+^ release events (Ca^2+^ blips and quarks) are often observed as precursors to puffs, suggesting that these low-amplitude Ca^2+^ release events trigger full-sized Ca^2+^ puffs and sparks (Rose et al., 2006). This is consistent with the observation that individual IP_3_Rs and RyRs are activated by cytosolic Ca^2+^, that is, small increases in [Ca^2+^] near these channels open these channels promotes further release of intracellular Ca^2+^, a process known as Ca^2+^-induced Ca^2+^ release (Bezprozvanny et al., 1991; Finch et al., 1991; Parker and Yao, 1996; Parker et al., 1996).

Although the activation mechanism of Ca^2+^ puffs and sparks is agreed upon, the mechanism by which puffs and sparks terminate is understood to a lesser degree and may vary in different physiological contexts (see Stern and Cheng (2004) for review). The short durations of most stochastic Ca^2+^ release events (10–200 ms) suggests that puff/spark termination is facilitated by a robust negative feedback mechanism (Cheng et al., 1993; Niggli and Shirokova, 2007). Because puff/sparks involve a finite number of channels, one possible termination mechanism is the simultaneous de-activation of all channels at a Ca^2+^ release site, a phenomenon referred to as stochastic attrition (DeRemigio and Smith, 2005; Groff and Smith, 2008a). Another possibility is that decreasing [Ca^2+^] in the SR/ER lumen reduces the driving force for Ca^2+^ release and/or the contribution of feed-through Ca^2+^-activation to channel closure (Huertas and Smith, 2007). The inhibitory role of cytosolic Ca^2+^-mediated inactivation of IP_3_Rs and RyRs is also thought to contribute to puff/spark termination (Fill, 2002; Stern and Cheng, 2004; Fraiman et al., 2006). Termination of stochastic Ca^2+^ release could also be mediated by state-dependent allosteric interactions between adjacent intercellular Ca^2+^ channels (Wiltgen et al., 2014), the redox state of IP_3_Rs and RyRs (Zima and Blatter, 2006; Hool and Corry, 2007), and luminal regulation mediated by calsequestrin or other ER/SR proteins (Györke et al., 2004).

Discrete-state continuous-time Markov chains (CTMCs) are often used to model the stochastic gating of plasma membrane and intercellular ion channels, including clusters of IP_3_Rs and RyRs collectively gating within CaRUs (Groff et al., 2010). These theoretical studies help clarify the factors that contribute to the generation and termination of Ca^2+^ puffs and sparks. Simulations show that moderately fast Ca^2+^ inactivation leads to puffs and sparks whose termination is facilitated by the recruitment of inactivated channels during the puff/spark event, while slow Ca^2+^ inactivation facilitates puff/spark termination due to stochastic attrition (DeRemigio and Smith, 2005; Groff and Smith, 2008b). Ca^2+^-mediated coupling of IP_3_Rs and RyRs also influences stochastic excitability of simulated CaRUs. The efficacy of this coupling is determined by the bulk ER/SR [Ca^2+^], the dynamics of luminal depletion, and the number, density and spatial arrangement of channels within a CaRU (Nguyen et al., 2005; DeRemigio and Smith, 2008).

In this paper, we present a Langevin formulation of the stochastic dynamics of Ca^2+^ release mediated by IP_3_Rs and RyRs that are instantaneously coupled through a local ‘domain’ Ca^2+^ concentration (a function of the number of open channels). The Langevin approach assumes the number of Ca^2+^ channels in individual CaRUs is large enough that the fraction of channels in different states can be treated as a continuous variable. Importantly, the computational efficiency of the Langevin approach is linear in the number of channel states and independent of the number of Ca^2+^ channels per CaRU. This is quite distinct from compositionally defined Markov chain models, in which the number of CaRU states is exponential in the number of channel states and polynomial in the number of channels per CaRU. For this reason, the Langevin approach may be preferred for extensive parameter studies, provided the Langevin model of stochastic Ca^2+^ release is a sufficiently good approximation to the corresponding Markov chain.

The remainder of this paper is organized as follows. Section 2 presents a continuous-time Markov chain model (and the corresponding Langevin formulation) of a CaRU composed of N three-state channels, each of which exhibits fast Ca^2+^ activation and slower Ca^2+^ inactivation. Section 3.1 uses the Langevin CaRU model to illustrate how the mechanism of spark termination depends on the rate of Ca^2+^ inactivation. By comparing statistics of simulated puff/sparks (amplitude, duration and inter-event interval) generated by both models, Section 3.2 demonstrates that the Langevin description of the collective gating of Ca^2+^ channels is indeed a good approximation to the corresponding Markov chain model when the number of Ca^2+^ channels per release site is in the physiological range. Section 3.3 uses Langevin simulations of stochastic Ca^2+^ release to preform an investigation of the correlations between successive puff/spark amplitudes, durations and inter-spark intervals and the dependence of these puff/spark statistics on the number of channels per release site and the kinetics of Ca^2+^-mediated inactivation of open channels.

## 2 Model formulation

### 2.1 Markov chain model of a Ca^2+^ release site

The stochastic gating of intracellular channels is often modeled by discrete-state continuous-time Markov chains. For example, the following state and transition diagram,

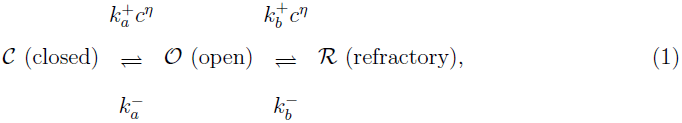

represents a minimal three-state channel that is both activated (𝓒 → 𝓞) and inactivated (𝓞 → 𝓡) by Ca^2+^ (Groff and Smith, 2008a). In this diagram, *c* is the local [Ca^2+^]; *η* is the cooperativity of Ca^2+^ binding; 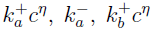 and 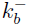 are transition rates with units of time^−1^; 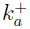 and 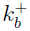 are association rate constants with units of concentration^−*η*^ time^−1^; and the dissociation constants for Ca^2+^ binding are 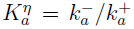 and 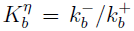. For simplicity, the cooperativity of Ca^2+^ binding is the same for the activation and inactivation processes (*η* = 2).

It is straightforward to construct a Ca^2+^ release unit (CaRU) model that includes an arbitrary number *N* of stochastically gating three-state channels. Because the channels are identical, such a model has (*N* + 2)(*N* + 1)/2 distinguishable states that may be enumerated as follows,

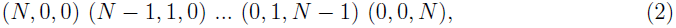

where each state takes the form (*N*_𝓒_, *N*_𝓞_, *N*_𝓡_) and *N*_𝓒_, *N*_𝓞_ and *N*_𝓡_ are the number of channels in closed, open and refractory states, respectively. For example, let us assume that when *N*_𝓞_ = *n*, the local [Ca^2+^] experienced by channels in the CaRU is given by

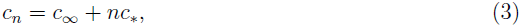

where *c*_∝_ is the bulk [Ca^2+^]. We will refer to the parameter *c*_*_ as the coupling strength, because this parameter determines the increment in local [Ca^2+^] that occurs when an individual Ca^2+^ channel opens. The transition rates for the compositionally defined Markov chain model with state space given by Eq. 2 are each the product of a transition rate of the single channel model (Eq. 1) and the number of channels that may make that transition. For example, in a release site composed of 20 channels, the transition rates out of the state (15,3,2) would be 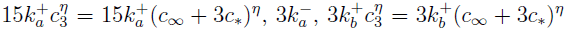, and 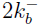 respectively, with destination states (14,4,2), (16,2,2), (15,2,3) and (15,4,1).

### 2.2 The Langevin description of a Ca^2+^ release site

A Langevin description of the CaRU is an alternative to the Markov chain model presented above (Gardiner, 1985; Dangerfield et al., 2012). The Langevin approach assumes that the number of channels in the CaRU is large enough so that the fraction of channels in each state can be treated as continuous randomly fluctuating variables that solve a stochastic differential equation (SDE) system. For example, the Langevin equation of a CaRU composed of *N* three-state channels (Eq. 1) is given by (Keizer, 1987)

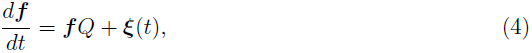

where ***f*** is a row vector of the fraction of channels in each state, ***f*** = (*f*_𝓒_, *f*_𝓞_, *f*_𝓡_), *Q* is the infinitesimal generator matrix (Q-matrix) given by

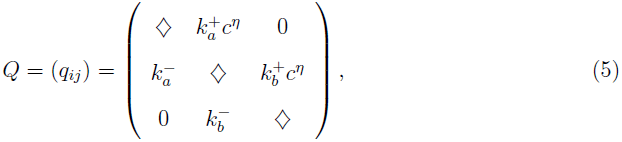

where the local [Ca^2+^] can be written as *c* = *c*_∝_ + *f*_𝓞_*c̄* with *c̄* = *N*_*c*_*__, the off-diagonal elements are transition rates (*q*_*ij*_ ≥ 0), and the diagonal elements (◇) are such that each row sums to zero, *q*_*ii*_ = – Σ_*j* ≠ *i*_ *q*_*ij*_ < 0. In Eq. 4, ***ξ***(*t*) = (*ξ*_𝓒_(*t*), *ξ*_𝓞_(*t*), *ξ*_𝓡_(*t*)) is a row vector of rapidly varying forcing functions with mean zero,

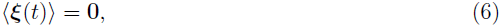

and two-time covariance,

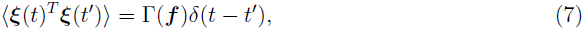

where Γ(***f***) = (*γ*_*ij*_) and

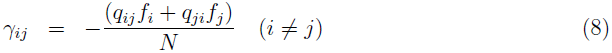

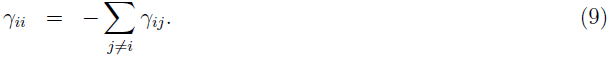

The Langevin model is simulated by integrating Eqs. 4–9 using a modification of the Euler-Maruyama method (Gillespie, 2000), appropriate for a stochastic ODE with dependent variables constrained to the unit interval, i.e., 0 ≤ *f_i_* ≤ 1 (see Wang et al. (2014) for details).

## 3 Results

The focus of this paper is a theoretical analysis of spark statistics such as puff/spark duration, amplitude and inter-event interval. We are specifically interested in the relationship between successive puff/spark amplitudes, whether puff/sparks and inter-event intervals are positively or negatively correlated, and how such puff/spark statistics depend on the single channel kinetics (e.g., Ca^2+^ inactivation rate). The Langevin approach to modeling CaRU dynamics facilities the large number of Monte Carlo simulations required for this analysis. Below we will first show representative Langevin simulation and illustrate how sequences of spark amplitudes, durations, and inter-event intervals are obtained from Langevin release site simulation (Section 3.1). Next we validate the Langevin release site model by comparing the statistics of simulated puff/sparks (amplitude, duration and inter-event interval) generated by the Langevin model and the corresponding Markov chain model (Section 3.2). This is followed by an analysis of correlations between successive puff/spark amplitudes, durations and inter-spark intervals, and how such puff/spark statistics depend on the number of channels per release site and the kinetics of Ca^2+^-mediated inactivation of open channels (Section 3.3).

### 3.1 Representative Langevin simulations

In prior work, Groff and Smith (2008a) found that Ca^2+^-dependent inactivation may facilitate puff/spark termination in two distinct ways depending on Ca^2+^-inactivation rates. Fig. 1A and B use the Langevin model (Eqs. 4–9) to illustrate these two different termination mechanisms. In Fig. 1A the number of inactivated channels (*N*_𝓡_, gray line) increases during each puff/spark event, and decreases during the inter-event intervals between puff/sparks.

**Fig. 1.**
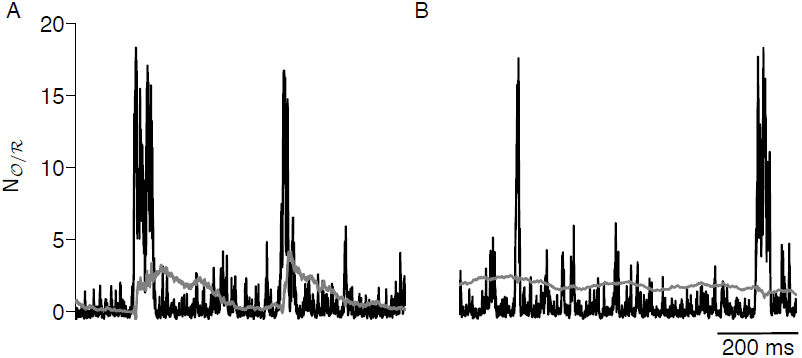
The number of open channels (*N*_𝓞_, black line) and refractory channels (*N*_𝓡_, gray line) during simulated Ca^2+^ puffs/sparks obtained by numerically integrating the Langevin model (Eqs. 4–9 with integration time step Δ*t* = 0.1 ms). Ca^2+^-inactivation/de-inactivation rates are 10-fold slower in B 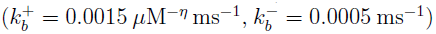 then A 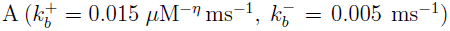. Other parameters: 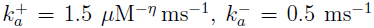, *c*_*_ = 0.06 *μ*M, *c*_∝_ = 0.05 *μ*M, *η* = 2, *K*_*a*_ = *K*_*b*_ = 0.58 *μ*M.

In this case, the Ca^2+^ inactivation rate is such that the accumulation of inactivated channels results in puff/spark termination. In Fig. 1B, the Ca^2+^ inactivation/de-inactivation rates are reduced by 10-fold compared with that of Fig. 1A. In this case the number of inactivated channels (*N*_𝓡_, gray line) is relatively constant; consequently, the CaRU composed of *N* three-state channels effectively reduces to a collection of *N* – *N*_𝓡_ two-state channels. In Fig. 1B, the puff/spark termination is due to stochastic attrition (Stern, 1992; Stern and Cheng, 2004), that is, the coincident de-activation (*N*_𝓞_ → *N*_𝓒_) of all channels in the CaRU that are not in the refractory state *N*_𝓡_ (Groff and Smith, 2008a).

Fig. 2A shows a Langevin simulation of the fraction of open channels, *f*_𝓞_, for a CaRU composed of 20 three-state Ca^2+^ channels. The duration of the *i*th Ca^2+^ release event (*D*_*i*_) is the time elapsed between the first channel opening (up arrows) and the last channel closing (down arrows) of each simulated spark. Because the fraction of open channels in the Langevin description is continuous (as opposed to discrete), the first/last channel opening is defined as *f*_𝓞_ crossing the threshold (1/*N*, dashed line) in the upward/downward direction (vertical arrows). The amplitude of ith Ca^2+^ release event (*A*_*i*_) is defined as the integrated area under *f*_𝓞_(*t*) during the release event (gray). The inter-event interval (*I*_*i*_) is the length of time between the (*i*–1)th and *i*th Ca^2+^ release events. Because in experimental studies many Ca^2+^ release events may be too small for detection, we specify an amplitude threshold, *A*_*θ*_, and only the events with greater amplitude (*A*_*i*_ ≥ *A*_*θ*_) are used in the calculation of spark statistics. For example, using an amplitude threshold of *A*_*θ*_ = 0.5 ms, only three of four events are of sufficient magnitude to be included in the sequence of puff/spark durations, amplitudes and inter-event intervals chosen for further analysis (Fig. 2B). If *A*_*θ*_ = 1 ms, only two of the four events are included (Fig. 2C).

**Fig. 2.**
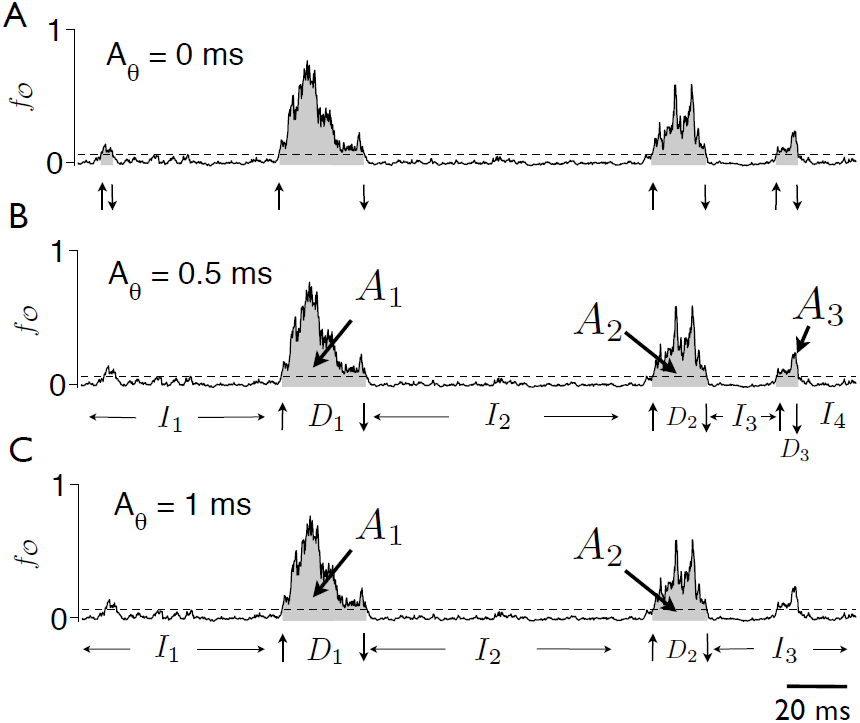
The puff/spark detectability threshold *A*_*θ*_ eliminates small Ca^2+^ release events from the correlation analysis of the sequence of simulated spark amplitudes, durations and interevent intervals. A: 20 three-state Ca^2+^ channels simulated using the Langevin approach (Eqs. 4–9). The fraction of open channels, *f*_𝓞_, is shown as a function of time. The dashed line denotes *f*_𝓞_ = 1/*N*, the threshold for identifying Ca^2+^ release events. Up and down arrows indicate crossings that define the beginning and ending of Ca^2+^ sparks. B: For an amplitude threshold *A*_¸_ = 0.5 ms, the first Ca^2+^ release event in A is discarded and three Ca^2+^ spark events are considered detectable. C: For Ca^2+^ = 1 ms, the first and last Ca^2+^ release event in A are discarded, and two Ca^2+^ spark events are detectable.

### 3.2 Validation of Langevin approach

Using CaRUs composed of *N* = 20 three-state channels and amplitude threshold of *A*_*θ*_ = 1 ms (lower dashed line) that filters out small events, Fig. 3A shows a strong linear relationship between spark amplitudes and durations in both Markov chain (∘) and Langevin (+) simulations. Using *A*_*θ*_ = 1 or 2 ms, Fig. 3B–D compares the cumulative distribution functions of spark amplitude (B), duration (C) and inter-event interval (D) for CaRUs composed of 20 (thin) and 60 (thick lines) three-state channels. In these simulations, the aggregate coupling strength *c̄* = *N*_*c*_*__ is fixed, as opposed to fixing the contribution to the local [Ca^2+^] made by a single open channel (*c*_*_). The spark duration and inter-event interval distributions move to the right as the number of channels increases (Fig. 3C–D), that is, for larger *N*, spark durations and inter-event intervals are typically longer. At the same time, increasing *N* leads to sparks that on average have smaller amplitudes (Fig. 3B). For high amplitude threshold *A*_*θ*_, the distributions move to the right because smaller events are filtered out. Most importantly, the agreement between Markov chain (black solid) and Langevin (gray dashed line) calculations of sparks statistics shown in Fig. 3B–D validates our use of Langevin approach for further analysis.

**Fig. 3.**
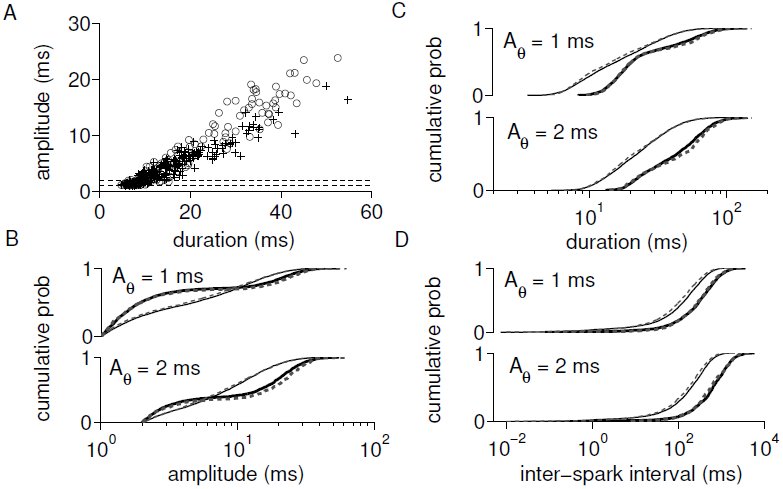
Spark duration, amplitude and inter-event interval statistics. A: Scatter plot of spark amplitudes vs. durations in Langevin (+) and Markov chain (∘) simulations shows a strong linear dependence (*N* = 20 channels) for *A*_*θ*_ = 1 ms and 2 ms (dash lines). B-D: The cumulative probability distributions of spark amplitude, duration and inter-event interval in Langevin (dashed) and Markov chain (solid lines) simulations for *A*_*θ*_ = 1 and 2 ms. Parameters as in Fig. 1A with *c̄* = 1.2 *μ*M and *N* = 20 or 60 (thin and thick lines, respectively).

### 3.3 Analysis of spark statistics

Our analysis of puff/sparks statistics begins with Fig. 4A which shows the Pearson correlation coefficient between successive puff/spark amplitudes, *ρA*_*n*_, *A*_*n* + 1_, when the standard parameters for Ca^2+^-inactivation and de-inactivation are used (as in Fig. 1A, where sparks terminate through the accumulation of inactivated channels). The correlation between successive puff/spark amplitudes is small but negative regardless of the amplitude threshold (*A*_*θ*_), indicating event-to-event alternation of puff/spark amplitude (small, large, small, large, etc.). The alternation in puff/spark amplitudes is most pronounced (i.e., the correlation is most negative) when *A*_*θ*_ is about 0.5 ms for *N* = 20 channels and 1 ms for 60 channels.

**Fig. 4.**
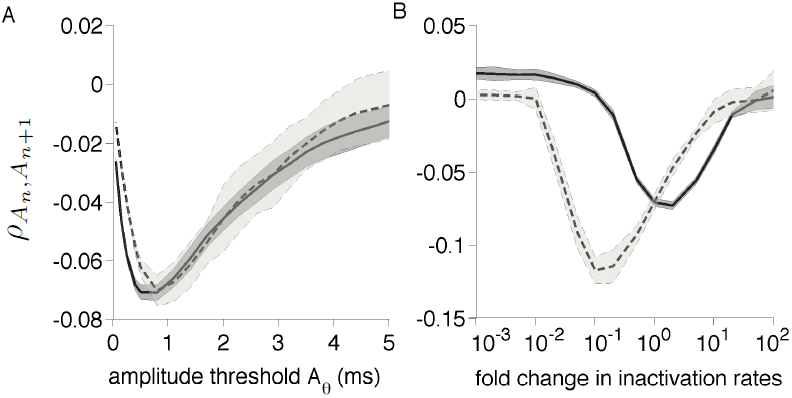
Correlation between successive puff/spark amplitude (*ρA*_*n*_, *A*_*n* + 1_). A: Correlation as a function of amplitude threshold *A*_*θ*_. B: The minimum correlation measured for a range of amplitude thresholds (*A*_*θ*_) as a function of inactivation/de-inactivation rates for fixed dissociation constant. In (A) and (B), the mean (thick curve) +/– one standard deviation (shaded region) over 10 simulations (each 8000 s) is shown. Parameters as in Fig. 1A with *N* = 20 (solid) and 60 (dashed lines) and *c̄* = 1.2 *μ*M.

Fig. 4B shows how this tendency toward alternating puff/spark amplitude depends on the rate of Ca^2+^-inactivation/de-inactivation (with dissociation constant *K*_*b*_ fixed). As a summary, we plot the minimum value of the correlation coefficient observed over a range of amplitude thresholds (0.05 ≤ *A*_*θ*_ ≤ 5 ms). Negative correlation between successive puff/spark amplitudes is observed for intermediate inactivation rates. This negative correlation occurs because large puff/sparks terminate with a relatively large fraction of inactivated channels (cf. Fig. 1A). Consequently, fewer channels are available to participate the next (small amplitude) Ca^2+^ release event. Conversely, small puff/sparks terminate with fewer Ca^2+^-inactivated channels, and the subsequent puff/spark amplitudes are thus likely to be larger. Negative amplitude-amplitude correlation is not observed when the inactivation rates are reduced or increased by 100-fold compared to the standard parameters. When the inactivation rate is slow enough that the number of refractory channels *N*_𝓡_ is essentially constant, the mechanism that may generate negative amplitude-amplitude correlation is no longer operative, because *A*_*n*_ does not affect *N*_𝓡_ at puff/spark termination. Similarly, when the inactivation rates are very fast, *A*_*n*_ can not influence *N*_𝓡_ at spark termination because *N*_𝓡_ is in quasistatic equilibrium with 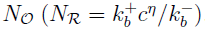. For both very slow and very fast inactivation rates, stochastic attrition is the mechanism of spark termination and spark amplitudes are less correlated. Interestingly, for a larger number of channels (*N* = 60), the most pronounced amplitude alternation is larger in magnitude (i.e., a stronger negative correlation) and occurs at slower inactivation rates than the *N* = 20 case.

Fig. 5 is similar in structure to Fig. 4, but focuses on the correlation between inter-event intervals and the subsequent puff/spark amplitudes, *ρI*_*n*_, *A*_*n*_, which are positively correlated regardless of amplitude threshold *A*_*θ*_ (Fig. 5A). Following a long inter-event interval, the channels that were inactivated at the end of the preceding puff/spark are more likely to be available for the subsequent Ca^2+^ release event. Consequently, the spark amplitudes following long quiescent periods tends to be larger than those that follow brief quiescent periods. Fig. 5B shows that the interval-amplitude correlation becomes very small for sufficiently slow or fast inactivation rates, and a larger correlation peak is observed for larger *N*. Similarly, Fig. 6A shows a small positive correlation between puff/spark amplitudes and the subsequent inter-event intervals (*ρA_n_*, *I*_*n*+1_). Fig. 6B shows that this positive amplitude-interval correlation becomes negligible for sufficiently fast or slow inactivation rates (similar to the interval-amplitude correlation of Fig. 5B).

**Fig. 5.**
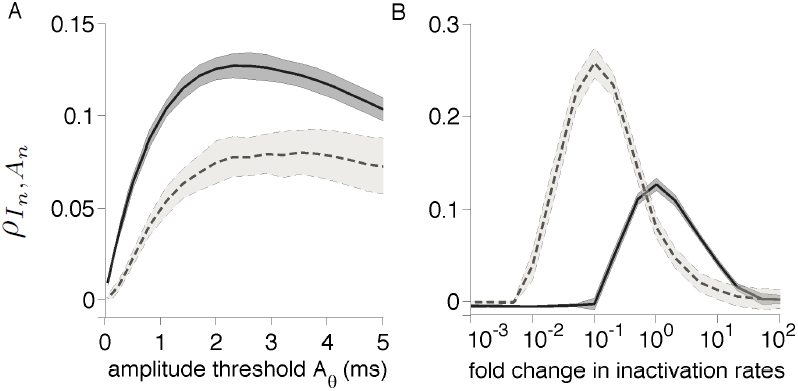
Correlation between inter-spark interval and subsequent puff/spark amplitude (*ρI*_*n*_, *A*_*n*_). A: Correlation as a function of amplitude threshold *A*_*θ*_. B: The maximum correlation measured for a range of amplitude thresholds (*A*_*θ*_) as a function of inactivation/deinactivation rates for fixed dissociation constant. See Fig. 4 legend.

**Fig. 6.**
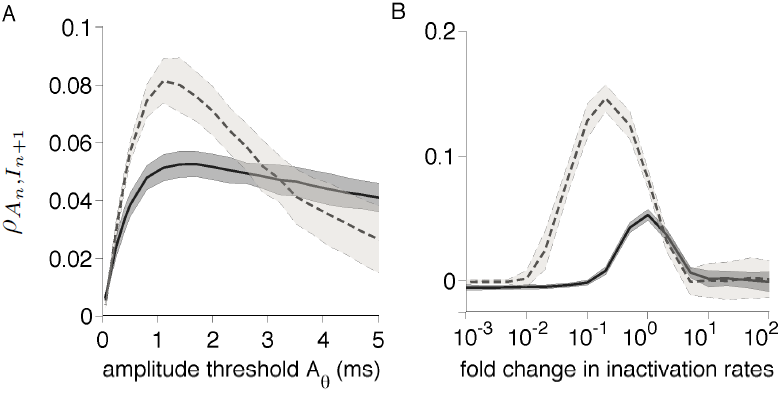
Correlation between puff/spark amplitude and subsequent inter-spark interval (*ρA*_*n*_, *I*_*n* + 1_). A: Correlation as a function of amplitude threshold *A*_*θ*_. B: The maximum correlation measured for a range of amplitude thresholds (*A*_*θ*_) as a function of inactivation/deinactivation rates for fixed dissociation constant. See Fig. 4 legend.

## 4 Discussion

This paper presents a Ca^2+^ release unit (CaRU) modeling approach based on a Langevin description of stochastic Ca^2+^ release. This Langevin model facilitates our investigation of correlations between successive puff/spark amplitudes and inter-spark intervals, and how such puff/spark statistics depend on the number of channels per release site and the kinetics of Ca^2+^-mediated inactivation of open channels. We find that when Ca^2+^ inactivation/deinactivation rates are intermediate—i.e., the termination of Ca^2+^ puff/sparks is caused by the recruitment of inactivated channels—the correlation between successive puff/spark amplitudes is negative, while the correlations between puff/spark amplitudes and the duration of the preceding or subsequent inter-spark interval are positive. These correlations are significantly reduced when inactivation/de-inactivation rates are extreme (slow or fast), that is, when puff/sparks terminate via stochastic attrition.

### 4.1 Comparison to experiment

Puff/spark amplitudes, durations and inter-event intervals have been extensively studied in recent years (Yao et al., 1995; Cheng et al., 1999; Rios et al., 2001; Shuai and Jung, 2002; Shen et al., 2004; Ullah and Jung, 2006). Positive correlations between spark amplitude and duration (Cheng et al., 1999) and rise time (Shkryl et al., 2012) have been observed. The rise time of spark fluorescence is interpreted as a proxy for the duration of Ca^2+^ release during the spark. Spark amplitude in our release site simulations is an increasing function of spark duration (i.e., positively correlated, with correlation coefficient of *ρ* = 0.95).

This positive correlation is not always observed in experimental (Wang et al., 2002) and theoretical work (Shuai and Jung, 2002). Shen et al. (2004) suggests that spark amplitude is independent of rise time, but is strongly and positively related to the mean or maximal rising rate. Their results show that spark rising time is negatively related to the mean rising rate, suggesting that the regulation of Ca^2+^ termination is a negative feedback and the strength of which is proportional to the ongoing release flux or the number of activated RyRs (Shen et al., 2004). In contrast to prior experimental results (Shkryl et al., 2012), the amplitude-rising time relationship is negative in the simulation work presented by Stern et al. (2013), because a long rise time implies a slower release of approximately the same amount of junctional SR Ca^2+^.

In our simulations, successive puff/spark amplitudes are negatively correlated, but only weakly (the peak is less than 0.15). This is consistent with experimental measurements of the correlation between successive puff amplitude were not statistically significant (Callamaras and Parker, 2000).

Inter-event intervals are determined both by recovery from a refractory state established by the preceding puff/spark, and a stochastic triggering which leads to an exponential distribution at longer intervals (Yao et al., 1995; Parker and Wier, 1997). The histograms of inter-puff interval measured at individual puff site show an initial increase of the inter-puff interval distribution, which is compatible with recovery from a negative feedback occurring during the puff (Thurley et al., 2011). In ventricular myocytes, Sobie et al. (2005) found that the relative amplitude of the second spark tends to be small when the spark-to-spark delay is short and larger as this delay increases. Moreover, Fraiman et al. (2006) observed that in *Xenopus* oocytes, puffs of large amplitudes tend to be followed by a long inter-puff time, and puffs that occur after a large inter-puff time are most likely large. One possible explanation of this positive interval-amplitude and amplitude-interval correlation is that high cytosolic [Ca^2+^] attained during a puff/spark inhibits channels within the CaRU, so that the amplitude and probability of occurrence of a subsequent puff recover with a long time course (Fraiman et al., 2006). Another possible mechanism is local Ca^2+^ depletion of ER lumen leading to decreased channel open probability (Fraiman and Dawson, 2004). Parker and Wier (1997) studied the relationship between the preceding inter-spark interval and the amplitude peak of the spark at the end of the interval and found no correlation. Our simulation exhibit a positive correlation between the preceding inter-spark interval and the amplitude of the spark at the end of the interval, but the maximum correlation observed is always less than 0.3 (Fig. 5B). The correlation between the spark amplitude and the subsequent inter-event interval is even weaker and the peak is less than 0.2 (Fig. 6B).

### 4.2 Comparison to prior theoretical work

Several types of modeling approaches based on microscopic kinetics of channels have been developed to study puff/spark statistics. For example, Ullah and Jung (2006) simplified the Sneyd-Dufour model (Sneyd and Dufour, 2002) and utilized it to study the spread of Ca^2+^ in the cytosol. They computed the correlation between puff amplitude and lifetime which is measured as the full width at half-maximal amplitude (FWHM). The predicted correlation of 0.31 indicates that puffs with larger amplitudes are likely to have longer lifetimes.

Ullah et al. (2012) presented a model of IP_3_R derived directly from single channel patch clamp data. Their results suggest that puff terminations is due to self-inhibition rather than ER Ca^2+^ depletion (unlike cardiac muscle, where local SR depletion is important for spark termination (Zima et al., 2008)).

Stern et al. (2013) utilized a simplified, deterministic model of cardiac myocyte couplon dynamics to show that spark metastability depends on the kinetic relationship of RyR gating and junctional SR refilling rates. They found that spark amplitudes is negatively correlated to rise time, in spite of the fact that positive correlation between amplitudes and rise time was observed in chemically skinned cat atrial myocytes (Shkryl et al., 2012).

Some prior work utilizing the Langevin formulation to investigate puff statistics has focused on a reduced Hodgkin-Huxley-like IP_3_ receptor model in which noise terms were added to the gating variable (Li and Rinzel, 1994; Shuai and Jung, 2002; Huang et al., 2011). By comparing Langevin and Markov chain simulations, they determined that Langevin approach yields more puffs with larger amplitudes, which leads to a drop-off of distribution at smaller amplitude; we did not observe this discrepancy in our Langevin simulation. Jung and coworkers also investigated the correlation between puff amplitude and lifetime and found that the correlation values are typically smaller than 0.3 (Shuai and Jung, 2002).

### 4.3 Advantages and limitations of the Langevin approach

While the Markov chain and Langevin approaches lead to similar results, the runtimes for Langevin simulations is often shorter. One expects Markov chain simulation runtimes to be proportional to the number of CaRU states, a quantity that is exponential in the number of distinct channel states. To see this, consider a CaRU composed of N identical M-state channels, the number of distinguishable CaRU states is given by (*N* + *M* – 1)!/[*N*!(*M* – 1)!]. For example, for 20, 60, and 100 identical three-state channels, there are 231, 1891, and 5151 distinguishable states, respectively. Conversely, Langevin simulation runtimes such as those presented in this study are independent of the number of channels (i.e., *N* is a model parameter) and proportional to the number of states *M*. Table A.1 illustrates this by comparing the simulation time of the Markov chain and the Langevin release site calculations shown here. The runtime of the Markov chain model increases significantly as the number of channels per release site *N* increases, while the simulation time of Langevin description is independent in *N*.

The Langevin formulation presented here is applicable and efficient when the number of channels per release site is large enough so that the fraction of channels in each state can be treated as a continuous variable. When this condition is not met, the use of Markov chain simulation or the slightly less-restrictive *τ*-leaping approach may be more appropriate (Gillespie, 2000).

This study has focus on correlations between spark statistics using relatively restrictive modeling assumptions, including a minimal three-state channel model and instantaneous coupling of channels. One important observation is that correlations between the amplitude, duration, and interval-event intervals of simulated Ca^2+^ puffs and sparks are strongly influenced by spark termination mechanism (i.e., Ca^2+^-dependent inactivation or stochastic attrition-like). Our formulation can be generalized to account for luminal depletion and/or regulation, both of which known to influence spark termination (Huertas and Smith, 2007), and finite system size effects (Weinberg and Smith, 2012).

## 5 ACKNOWLEDGMENTS

The work was supported in part by National Science Foundation Grant DMS 1121606 to GDS and the Biomathematics Initiative at The College of William & Mary.

## Appendix A Runtime of the Markov chain and Langevin model

**Table A.1.** Simulation time of a CaRU composed of *N* three-state channels, where *N* is 20, 60 and 100 respectively. Time step Δ*t* is 0.1 or 0.01 ms. The reported simulation times are the average of 10 100 s trials. Parameters as in Fig. 1A.

| Model | Simulation time (s) | Standard deviation (s) |
| --- | --- | --- |
| Markov chain ($N = 20$) | 11.78 | 0.56 |
| Markov chain ($N = 60$) | 148.86 | 2.37 |
| Markov chain ($N = 100$) | 416.80 | 4.26 |
| Langevin ($N = 20, \Delta t = 0.1$ ms) | 4.38 | 0.32 |
| Langevin ($N = 60, \Delta t = 0.1$ ms) | 4.28 | 0.35 |
| Langevin ($N = 100, \Delta t = 0.1$ ms) | 4.30 | 0.36 |
| Langevin ($N = 20, \Delta t = 0.01$ ms) | 43.98 | 3.14 |
| Langevin ($N = 60, \Delta t = 0.01$ ms) | 43.26 | 3.89 |
| Langevin ($N = 100, \Delta t = 0.01$ ms) | 44.47 | 3.52 |

